# Spatiotemporal regulation of GIPR signaling impacts glucose homeostasis as revealed in studies of a common GIPR variant

**DOI:** 10.1101/2020.05.12.091025

**Authors:** Lucie Yammine, Belén Picatoste, Nazish Abdullah, Rosemary A. Leahey, Emma F. Johnson, Nicolás Gómez-Banoy, Carolina Rosselot, Jennifer Wen, Tahmina Hossain, Marcus D. Goncalves, James C. Lo, Adolfo Garcia-Ocaña, Timothy E. McGraw

## Abstract

Glucose-dependent insulinotropic polypeptide (**GIP**) has a role in controlling postprandial metabolic tone. In humans, a GIP receptor (**GIPR**) variant (Q354, rs1800437) is associated with a lower body mass index (**BMI**) and increased risk for Type 2 Diabetes. To isolate the contribution of GIPR in metabolic control, we generated a mouse model of the GIPR-Q354 variant (GIPR-Q350 mice). Female GIPR-Q350 mice are leaner than littermate controls, and male GIPR-Q350 mice are resistant to diet-induced obesity, in line with the association of the variant with reduced BMI in humans. GIPR-Q350 mice of both sexes are more glucose tolerant and exhibit an increased sensitivity to GIP. Postprandial GIP levels are reduced in GIPR-Q350 mice, revealing feedback regulation that balances the increased sensitivity of GIP target tissues to secretion of GIP from intestinal endocrine cells. The increased GIP sensitivity is recapitulated *ex vivo* during glucose stimulated insulin secretion assays in islets. Generation of cAMP in islets downstream of GIPR activation is not affected by the Q354 substitution. However, post-activation traffic of GIPR-Q354 variant in β-cells is altered, characterized by enhanced intracellular dwell time and increased localization to the Trans-Golgi Network (**TGN**). Consequently, our data link altered intracellular traffic of the GIPR-Q354 variant with GIP control of metabolism. We propose that this change in spatiotemporal signaling underlies the physiologic effects of GIPR-Q350/4 and GIPR-E350/4 in mice and humans. These findings contribute to a more complete understanding of the impact of GIPR-Q354 variant on glucose homeostasis that could perhaps be leveraged to enhance pharmacologic targeting of GIPR for the treatment of metabolic disease.

## Introduction

The glucose-dependent insulinotropic peptide (**GIP**) is a hormone secreted by the K cells of the intestinal epithelium in response to the caloric content of the chyme [1; 2]. GIP and another gut-derived hormone, Glucagon-like Peptide 1 (**GLP-1**), which together are referred to as incretin hormones, contribute significantly to control of postprandial metabolic tone [3–5]. Insulin secreting pancreatic β−cells are a target of incretin hormone action, with the incretins enhancing glucose-stimulated insulin secretion [6; 7]. The GIP receptor (**GIPR**) is more widely expressed than the GLP-1 receptor [8], and exerts extra-pancreatic roles in neurogenesis [9], fat accumulation in the adipose tissue [10] and bone formation [11].

Incretin mimetic drugs have rapidly gained interest in the treatment of obesity and type 2 Diabetes (**T2D**). While earlier studies had focused on GLP-1R agonism and dipeptidyl peptidase-4 inhibition [12; 13], GIPR targeting has attracted attention in more recent advancements [14–16]. In fact, GIPR/GLP-1R dual-agonist drugs have demonstrated high efficacy in the treatment of obesity and T2D in clinical studies, gaining a recent FDA approval [17; 18].

GIPR is a class B G protein-coupled receptor (**GPCR**) linked to adenylate cyclase activation via Gs [19]. Elevation of 3′-5′-cyclic adenosine monophosphate (**cAMP**) downstream of GIPR activation enhances glucose-stimulated insulin secretion [20]. Although GPCR signal activation occurs at the plasma membrane following ligand binding, the post activation traffic of GPCRs is crucial for sculpting response to receptor activation [21]. GIPR trafficking has been analyzed in a variety of cell types [22–24]. In our previous studies of GIPR in cultured adipocytes, we have shown that GIPR is constitutively internalized and recycled back to the plasma membrane, independent of ligand stimulation [25]. GIP stimulation slows recycling of the receptor without affecting GIPR internalization [25; 26]. The slowed recycling is achieved by diverting the active receptor from the endosomal recycling pathway to the slower Trans Golgi Network (**TGN**) recycling route [26]. The reduced recycling rate of active GIPR induces a transient downregulation of GIPR from the plasma membrane that is reversed when GIP-stimulation is terminated [25].

Genome-wide association studies have identified several naturally occurring variants of GIPR. GIPR-Q354 (SNP rs1800437), a substitution of glutamine for glutamic acid at position 354 of human GIPR [27–29], is a variant with a relatively high allele frequency of 0.2 in European descents. Homozygosity for GIPR-Q354 is associated with lower BMI in different meta-analyses, whereas the predominant GIPR form (GIPR-E354) has been associated with a higher susceptibility to obesity [28; 30; 31]. Individuals homozygous for the GIPR-Q354 variant have a reduced C-peptide excursion in response to an oral glucose tolerance test, as well as a reduced GIP and insulin concentrations, documenting effects of the GIPR-Q354 variant on β cell biology [32].

The substitution of glutamine for glutamic acid at position 354 of the GIPR variant does not affect the affinity for GIP nor GIP-stimulated increase in cAMP, demonstrating that the substitution does not affect the most receptor proximal aspects of GIP-stimulated GIPR signaling [25; 26; 33; 34]. However, we have demonstrated post-activation trafficking differences between GIPR-E354 and GIPR-Q354. In adipocytes, the GIPR-Q354 variant undergoes enhanced GIP-stimulated downregulation from the plasma membrane coupled with an enhanced trafficking of GIPR to the TGN [25; 26]. Consequences of the altered post-activation trafficking of GIPR-Q354 are an increased localization of active GIPR-Q354 in the TGN and a slower repopulation of plasma membrane GIPR-Q354 following termination of GIP stimulation, thereby resulting in an enhanced and prolonged downregulation of GIPR-Q354 relative to GIPR-E354 [25].

In this study we generated a mouse model of the human GIPR-Q354 variant, in which glutamine is substituted for glutamic acid at position 350 of the mouse GIPR (GIPR-Q350, equivalent to human GIPR-Q354). We reveal a critical role for this amino acid at position 354 of the mouse GIPR in the biology of β−cells. Homozygous GIPR-Q350 female mice gain less weight on a normal chow diet while males are resistant to a diet-induced obesity. GIPR-Q350 mice of both sexes have a significant improvement in glucose tolerance, as compared to littermate WT C57BL/6J mice (GIPR-E354). GIPR-350 mice are also significantly more sensitive to GIP in a glucose tolerance test. This establishes an impact of GIPR-Q350 on whole body glucose metabolism. GIPR-Q350 islets have increased glucose-stimulated insulin secretion and an increased response to GIP stimulation, consistent with the differences between genotypes in the control of whole-body glucose homeostasis.

## Results

### GIPR rs1800437-C variant is associated with lower body mass index

To broadly investigate the phenotypes associated with the GIPR-Q354 single nucleotide polymorphism (**SNP**) (rs1800437-C allele), we analyzed Phenome-wide association (**PheWAS**) of GIPR-Q354 for metabolism relevant traits (**Figure 1**). These results were extracted from the Type 2 Diabetes Knowledge Portal database (https://t2d.hugeamp.org/) and are the meta-analysis of GWAS studies that had associated GIPR-Q354 SNP with different phenotypes. For our purposes, Glycemic and Anthropometric traits were of interest, and we excluded phenotypes that did not reach a significance of *p* < 5 x 10^−8^. Taking into consideration the effective sample size and *p* value, we found the strongest association of the GIPR-Q354 variant with a lower BMI (*p* = 2.23 x 10^−111^, *b* = −0.0256, *n* = 4,061,430). In support of this finding, the rs1800437-C allele was independently associated with reduced waist circumference (*p* = 3.68 x 10^−10^, *b* = −0.0198, *n* = 1,218,430) and waist-hip ratio (*p* = 4.31 x 10^−12^, *b* = −0.0082, *n* = 2,448,960). In addition, GIPR-Q354 was associated with lower fasting blood glucose adjusted for BMI (*p* = 2.28 x 10^−8^, *b* = −0.0099, *n* = 359,776). Paradoxically, despite these links with phenotypes beneficial to a healthy metabolism, GIPR-Q354 was positively associated with elevated HbA1c (*p* = 1.70 x 10^−11^, *b* = 0.0033, *n* = 750,799), and quite strongly with T2D adjusted for BMI (*p* = 2.53 x 10^−17^, *b* = 0.0644, *n* = 351,558), and the two-hour glucose level during an oral glucose tolerance test (**O-GTT**) adjusted for BMI (*p* = 8.09 x 10^−27^, *b* = 0.0977, *n* = 76,538). Although these associations are generated using an additive model comprising heterozygosity for the rs1800437-C allele, they clearly identify a link between GIPR-Q354 and altered metabolism. To better understand the different impacts of GIPR-Q354 and GIPR-E354 on metabolism, it is necessary to study these variants in an isogeneic background. To accomplish this objective, we used CRISPR-CAS9 editing to generate mouse models of GIPR-Q354 and GIPR-E354.

**Figure 1:**
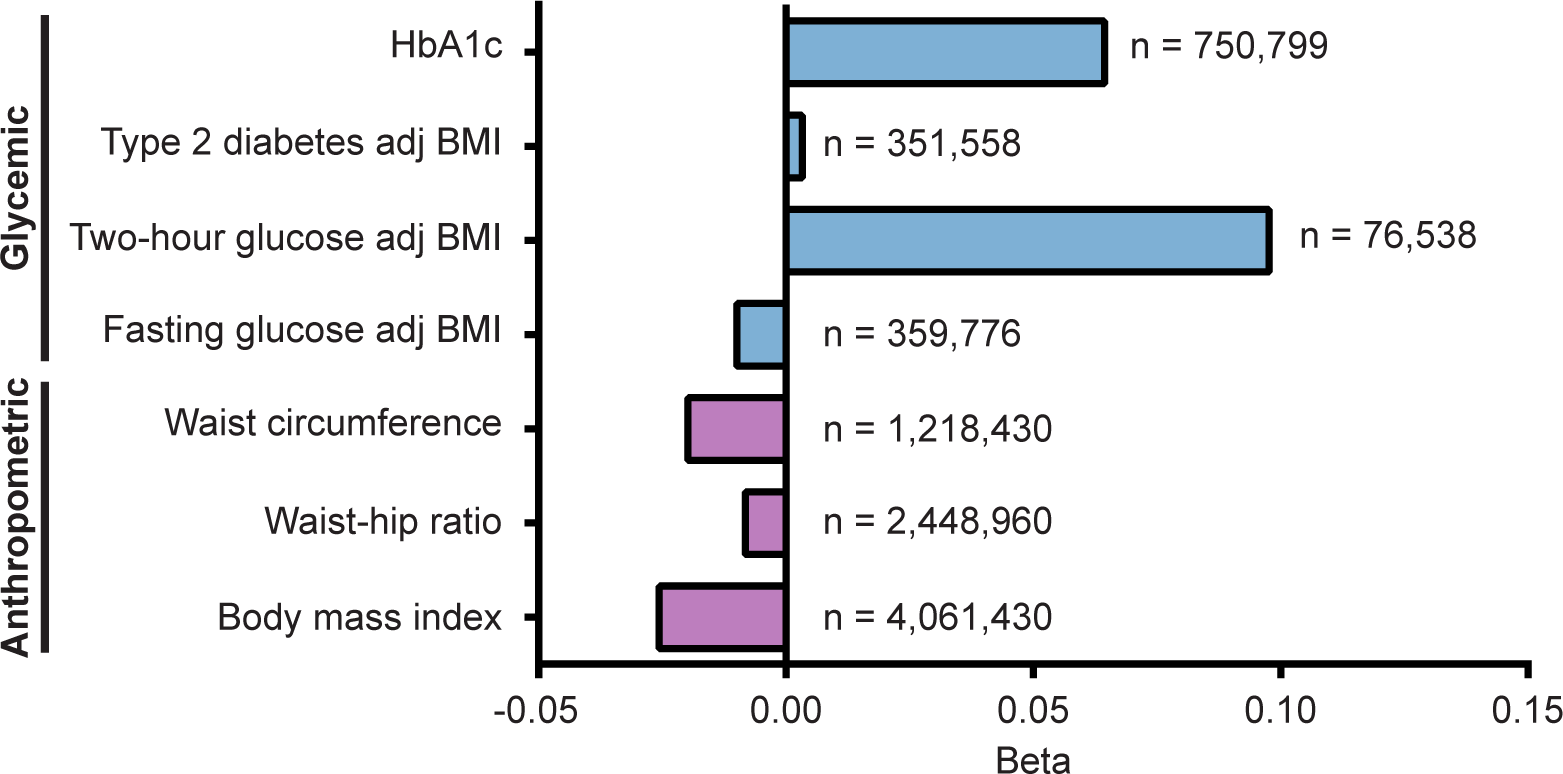
PheWAS shows GIPR-Q354 (rs1800437) decreased BMI phenotype. Graph shows significant (p < 5 x 10^−8^) associations of GIPR variant rs1800437-C for Glycemic and Anthropometric traits.

### Generation of GIPR-Q350 mice

Position 350 of the mouse GIPR is equivalent to position 354 of human GIPR. C57BL/6J mice have a glutamic acid at position 350 (**GIPR-E350**), and therefore are equivalent to the most common human GIPR protein sequence. To probe whether the differences in post-activation trafficking between GIPR-E354 and the GIPR-Q354 variant, discovered in our previous studies of cultured cells [25; 26], impacts GIP control of metabolic tone, we used CRISPR-CAS9 technology to generate a GIPR-Q350 variant in C57BL/6J mice. The genotype of the GIPR-E350 and GIPR-Q350 animals was confirmed by Sanger sequencing and PCR on genomic DNA using primers specific for the indel (**Fig. S1A**). GIPR mRNA expression was not altered in GIPR-Q350 mice, as compared to their respective GIPR-E350 controls (**Fig. S1B,C**), and the mice were born in the expected Mendelian frequencies.

### GIPR-Q350 mice are leaner than GIPR-E350 mice

When fed a normal chow diet, GIPR-Q350 female mice had a significantly lower body weight while maintaining comparable two-hour fasted blood glucose levels, as compared to age-matched GIPR-E350 females (**Figure 2A, B**). The lower weight of GIPR-Q350 female mice is consistent with the association of GIPR-Q354 with the lower body mass index in humans revealed by PheWAS analyses (**Figure 1**).

**Figure 2:**
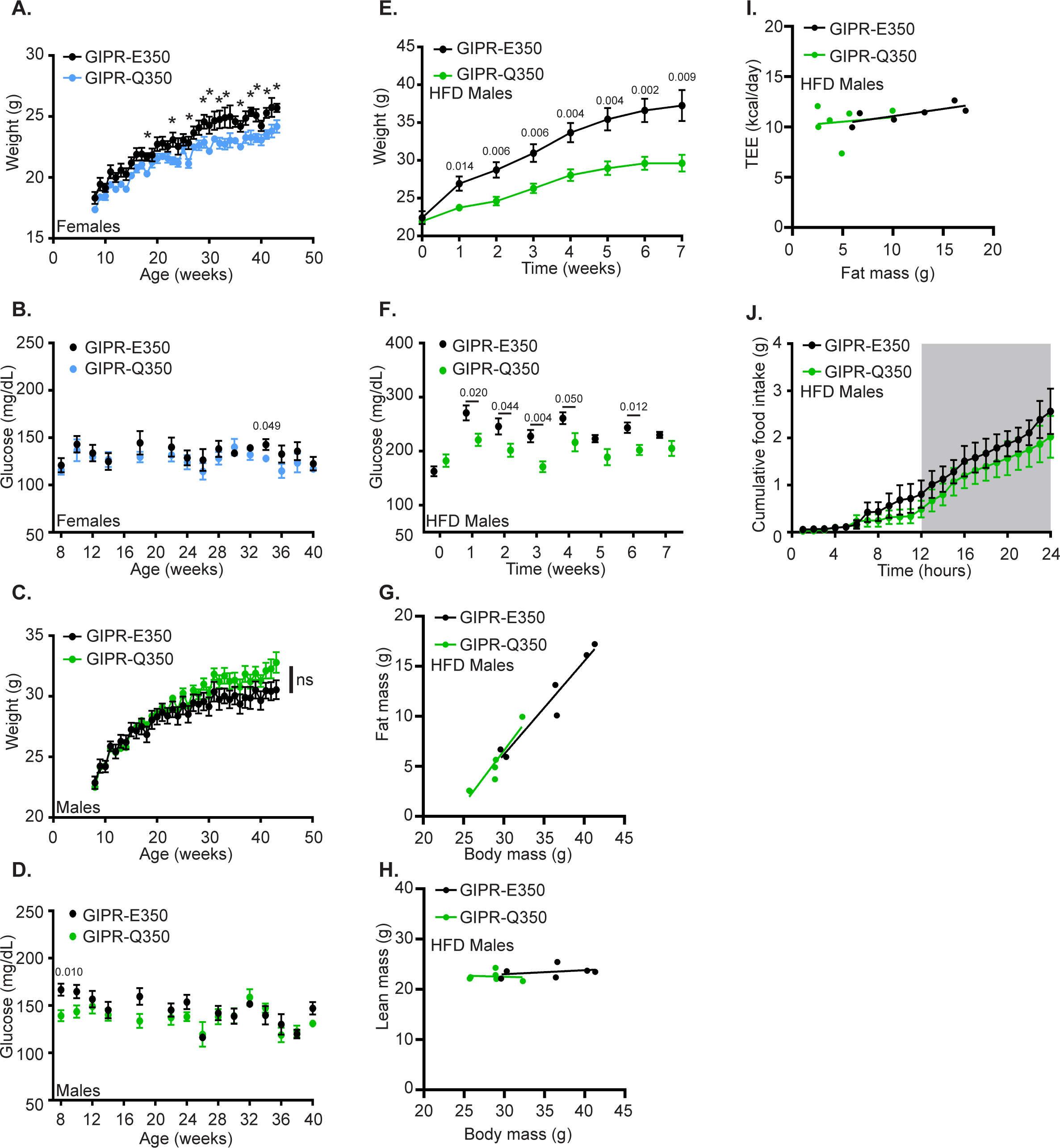
GIPR-Q350 mice recapitulate human GIPR-Q354 body weight increase. **A-D.** 8 weeks old GIPR-E50 and GIPR-Q350 female **(A&B)** and male **(C&D)** mice were monitored for their weights **(A&C)** and 2 h fasted blood glucose levels **(B&D)** over time. (n = 6 mice/genotype). **E&F.** 8 to 9 weeks old GIPR-E350 and GIPR-Q350 male mice were fed a 60% fat diet and their weights **(E)** and 2 h fasted blood glucose levels **(F)** were monitored over time. **G&H.** Fat mass **(G)** and Lean mass **(H)** compared to body mass as determined by MRI after 8 weeks on the HFD. **I.** Total daily energy expenditure (TEE) in relation to fat mass in HFD-fed males. **J.** Indirect food intake measurements of GIPR-E350 and GIPR-Q350 HFD-fed males over one light/dark cycle (24 h). (n = 6 mice/genotype). Data are mean ±SEM. Two tailed unpaired *t*-tests for A-F.

When fed a normal chow diet, both genotypes of male mice had comparable body weight gain over time, and there were no differences in two-hour fasted blood glucose levels between the genotypes (**Figures 2C,D**). However, a difference in body weights was revealed when mice were fed a high fat diet (HFD, 60% fat, 20% proteins, 20% carbohydrates) to mimic diet-induced obesity. The GIPR-Q350 mice gained less weight and maintained lower blood glucose levels over time, as compared to GIPR-E350 mice (**Figures 2E,F**). The reduced body weight was due to lower fat mass without a change in lean mass of HFD-fed GIPR-Q350 mice compared to GIPR-E350 mice (**Figure 2G,H**). Of note, the GIPR-Q350 male mice fed a HFD had similar body fat (16.6%) as similarly aged C57BL/6 mice (GIPR-E350) mice fed a normal chow diet [35; 36]. This indicates a near complete blunting of weight gain in GIPR-Q350 male mice on HFD. To gain insight into how the GIPR-Q350 mice maintained a lower body weight on HFD, we measured energy expenditure and food intake in single-housed mice 8 weeks after starting the HFD. There was a linear relationship between total daily energy expenditure (TEE) and fat mass (**Figure 2I**) with no significant difference between GIPR-E350 and GIPR-Q350 mice when TEE was adjusted for body weight, lean mass, or fat mass by ANCOVA (*p*>0.05). There was a trend toward lower food intake in GIPR-Q350 mice, as compared to GIPR-E350 mice (**Figure 2J**), which may explain the differences in weight gain, body weight, and fat mass. Male GIPR-Q350 mice also displayed better glucose tolerance and less hyperinsulinemia after an O-GTT as compared to the GIPR-E350 males after 17 weeks of HFD feeding (**Fig. S2A-C**). These results are in line with the anthropometric traits identified in human PheWAS (**Figure 1**).

### GIPR-Q350 mice are more glucose tolerant and sensitive to GIP

GIPR-Q350 female mice were more glucose tolerant than weight-matched GIPR-E350 mice in an O-GTT (**Figure 3A, Fig. S3A**). Unexpectedly, the enhanced glucose tolerance was not associated with post-stimulation differences in plasma insulin amounts between the female genotypes (**Figure 3B**); however, there were significant reductions in plasma GIP during the O-GTT in GIPR-Q350 mice (**Figure 3C**). Male GIPR-Q350 mice trended towards increased glucose tolerance in an O-GTT (**Figure 3D**). The difference in area under the glucose curve achieved statistical significance (p < 0.05) when assessed over multiple cohorts (n=4) (**Fig. S3B**). As was the case for female mice, there were no differences in plasma insulin between genotypes of male mice during the O-GTT, whereas plasma GIP was reduced in GIPR-Q350 male mice (**Figures 3E,F**). In both sexes, plasma GLP-1 levels were not affected by the GIPR-Q350 variant during the O-GTT (**Figure S3C,D**), and there were no significant differences in whole body insulin sensitivity between the two genotypes of mice of both sexes when measured by intraperitoneal insulin tolerance test (**IP-ITT**) with or without injection of exogenous GIP (**Fig. S3E-H**).

**Figure 3:**
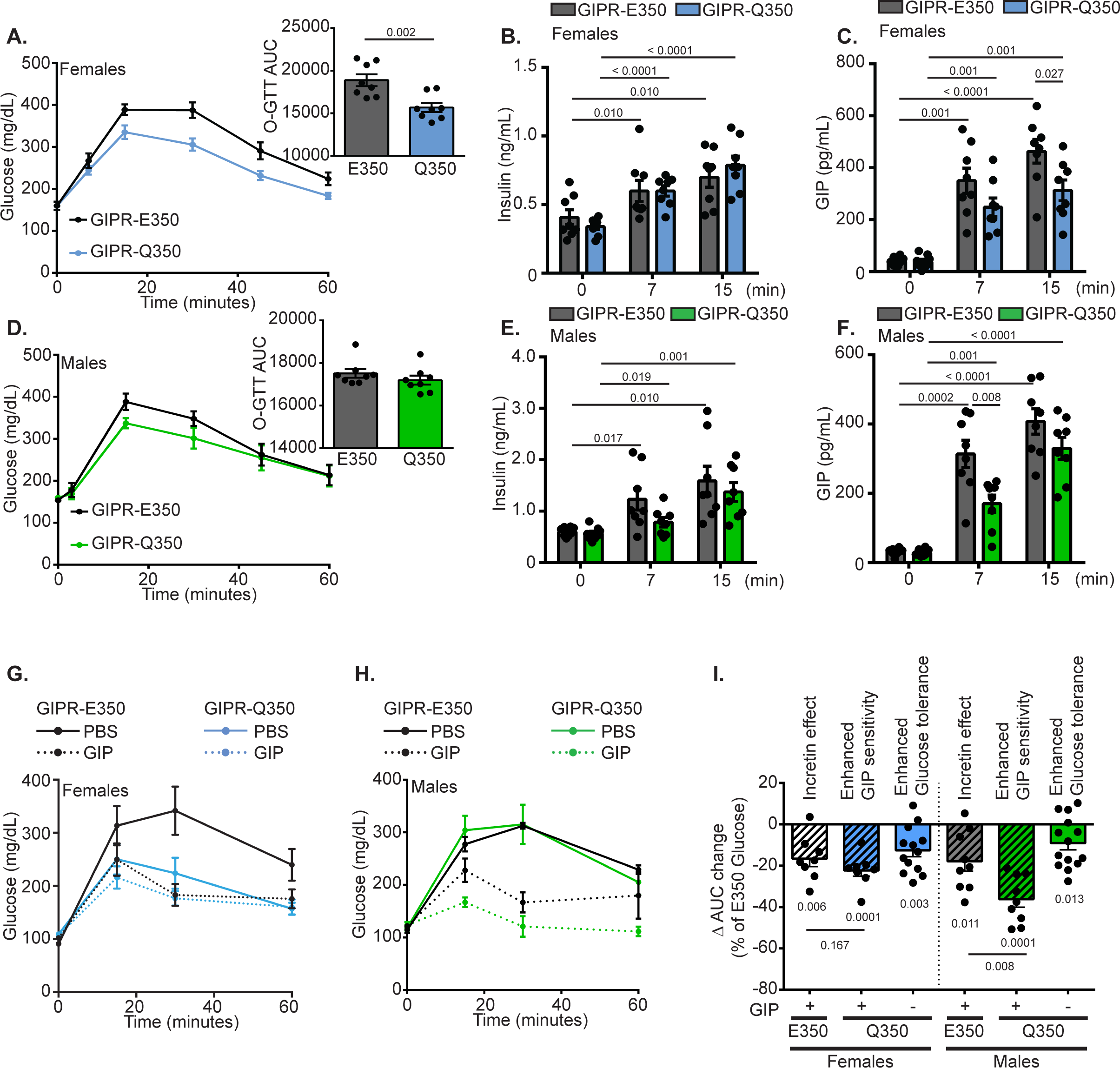
GIPR-Q350 mice are more glucose tolerant than GIPR-E350 mice. **A.** Blood glucose excursion and area under the curve (AUC, insert) over time after an oral glucose tolerance test (OGTT, 2 g/kg) in 6 h fasted GIPR-E350 and GIPR-Q350 females. (n = 8 mice/genotype). **B&C.** Plasma levels of Insulin **(B)** and GIP **(C)** before and at the indicated times after glucose administration (2 g/kg). (n = 8 mice/genotype). **D.** Blood glucose excursion and area under the curve (AUC, insert) over time after an oral glucose tolerance test (OGTT, 2 g/kg) in 6 h fasted GIPR-E350 and GIPR-Q350 males. (n = 8 mice/genotype). **E&F.** Plasma levels of Insulin **(E)** and GIP **(F)** before and at the indicated times after glucose administration (2 g/kg). (n = 8 mice/genotype). **G-I.** Blood glucose excursion **(G&H)** and delta area under the curve change (AUC, **I**) over time after an intraperitoneal glucose tolerance test (2 g/kg) supplemented or not with GIP (20 μmol/kg) normalized to GIPR-E350 glucose injection only condition in 16 h fasted GIPR-E350 and GIPR-Q350 females. (n = 8-14 independent experiments). Data are mean ±SEM. Two tailed unpaired *t*-tests for A, C, F. Two tailed paired *t*-tests for B, C, E, F, I.

GIP levels contribute to the amount of insulin secreted from β−cells following oral nutrient challenge (the incretin effect). Therefore, an unchanged level of insulin in context of lower plasma GIP suggests the GIPR-Q350 mice of both sexes have increased sensitivity to GIP. To isolate the effect of GIP on glucose stimulated insulin secretion (**GSIS**), we performed an IP-GTT in which we co-administered GIP with glucose injection (**Figure 3G-I**). This approach bypasses stimulation of endogenous GIP (and GLP-1) secretion, thereby isolating the incretin effect on glucose tolerance to that of the injected GIP. As expected, in IP-GTT, co-stimulation with GIP and glucose resulted in a better glucose tolerance in GIPR-E350 mice as compared to glucose alone (**Figure 3G-I**). The incretin effect of GIP in GIPR-E350 mice reduced the area under the curve (**AUC**) of the IP-GTT 16.6 ± 10.1 % (females) and 17.9 ± 13.6 % (males) compared to glucose alone (**Figure 3I**). The GIP incretin effect was larger in GIPR-Q350 mice, with co-injection of GIP and glucose resulting in significantly greater reductions in AUCs as compared to the GIP incretin effect in GIPR-E350 mice: 22.3 ± 7.9 % and 36.2 ± 10.8% for females and males, respectively (**Figures 3I**). This more robust incretin effect demonstrates increased GIP sensitivity of GIPR-Q350 β-cells in both sexes of mice. In addition, when averaged across multiple cohorts of mice, the IP-GTT AUC GIPR-Q350 mice was reduced by 12.6 ± 10.8 %, n = 5 cohorts for females and 9.1 ± 11.8%, n = 5 cohorts for males, which further confirms the improved glucose tolerance in GIPR-Q350 mice (**Figure 3I**).

To confirm the increase in the GIP incretin effect on insulin secretion in GIPR-Q350 mice, we dosed mice with increasing amounts of [D-Ala^2^]-GIP, a long-lasting GIP analog [37], in the setting of an IP-GTT (**Figure 4**). In female GIPR-E350 mice, only the 1 µg dose significantly reduced AUC as compared to vehicle, whereas both 0.5 and 1.0 µg doses were effective in the GIPR-Q350 mice (**Figure 4A-C**). In addition, the incretin effect of 1 µg [D-Ala^2^]-GIP was significantly stronger (lower AUC) in GIPR-Q350 females than in GIPR-E350 mice (**Figure 4C**). Similarly, in male GIPR-E350 mice, there was a trend toward a better GIP incretin effect with increasing doses, although it only reached significance at the 1 µg [D-Ala^2^]-GIP (**Figure 4E-G**). In the male GIPR-Q350 mice, both 0.5 and 1 µg doses significantly reduced AUC as compared to vehicle, and at 1 µg the incretin effect of [D-Ala^2^]-GIP in GIPR-Q350 males was enhanced compared to the GIPR-E350. In aggregate, these results confirm hypersensitivity of GIPR-Q350 mice to GIP.

**Figure 4:**
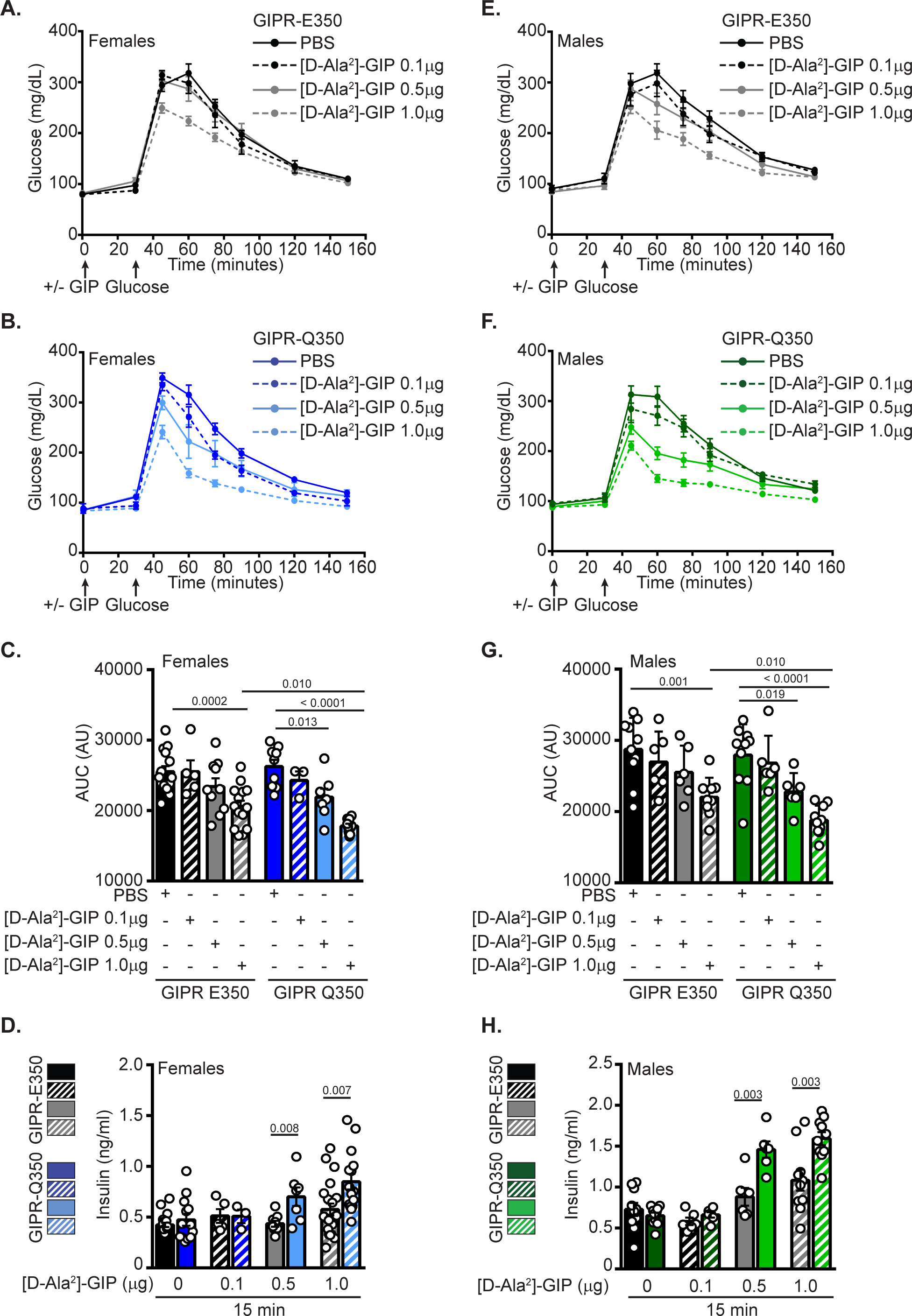
GIPR-Q350 mice are more sensitive to GIP than GIPR-E350 mice. **A-B.** Blood glucose excursion **(A&B)** and area under the curve **(C)** after an intraperitoneal glucose challenge (2 g/kg of body weight) assessed 30 min after an i.p. injection of either vehicle (PBS) or different doses of [D-Ala^2^]-GIP (0.1 μg, 0.5 μg, 1 μg) in 16 h fasted GIPR-E350 **(A)** and GIPR-Q350 **(B)** female mice. **D.** Plasma levels of Insulin at 15 minutes after IP-GTT (n = 7 – 16 mice/genotype). **E-G.** Blood glucose excursion **(E&F)** and area under the curve **(G)** after an intraperitoneal glucose challenge (2 g/kg of body weight) assessed 30 min after an i.p. injection of either vehicle (PBS) or different doses of [D-Ala^2^]-GIP (0.1 μg, 0.5 μg, 1 μg) in 16 h fasted GIPR-E350 **(E)** and GIPR-Q350 **(F)** male mice. **H.** Plasma levels of Insulin at 15 minutes after IP-GTT (n = 6 – 10 mice/genotype). Two tailed unpaired *t*-tests for C, D, G, H.

The improvement in IP-GTT AUC with 0.5 and 1.0 µg [D-Ala^2^]-GIP administration coincided with higher circulating insulin levels at the 15 min time point in GIPR-Q350 mice, as compared to GIPR-E350 mice (**Figures 4D,H**). There were no differences between genotypes in circulating insulin levels after an overnight fast (**Fig S4A,B**). The increased sensitivity of GIPR-Q350 mice to GIP did not affect response to GLP-1 of either sex (**Fig S4C,D**). Together, these data suggest that GIPR-Q350 β cells have increased sensitivity to GIP.

### Enhanced GSIS and incretin response in GIPR-Q350 islets *ex vivo*

Pancreatic β-cells are a primary target of GIP, and the enhanced glucose tolerance and increased insulin secretion suggest that the β-cell response to GIP is affected in GIPR-Q350 mice. We found no prominent differences in the cellular organization of the islets, with the insulin expressing β-cells segregated to the interior of islets and glucagon expressing α cells to the periphery (**Figure 5A**). Furthermore, there were no differences in pancreas weights or β-cell mass between genotypes (**Fig. S5A, B**). There was comparable expression (mRNA) of *Gipr*, *Glut2, Glp1r* and *Glucokinase* in both genotypes (**Fig. S5C,D**).

**Figure 5:**
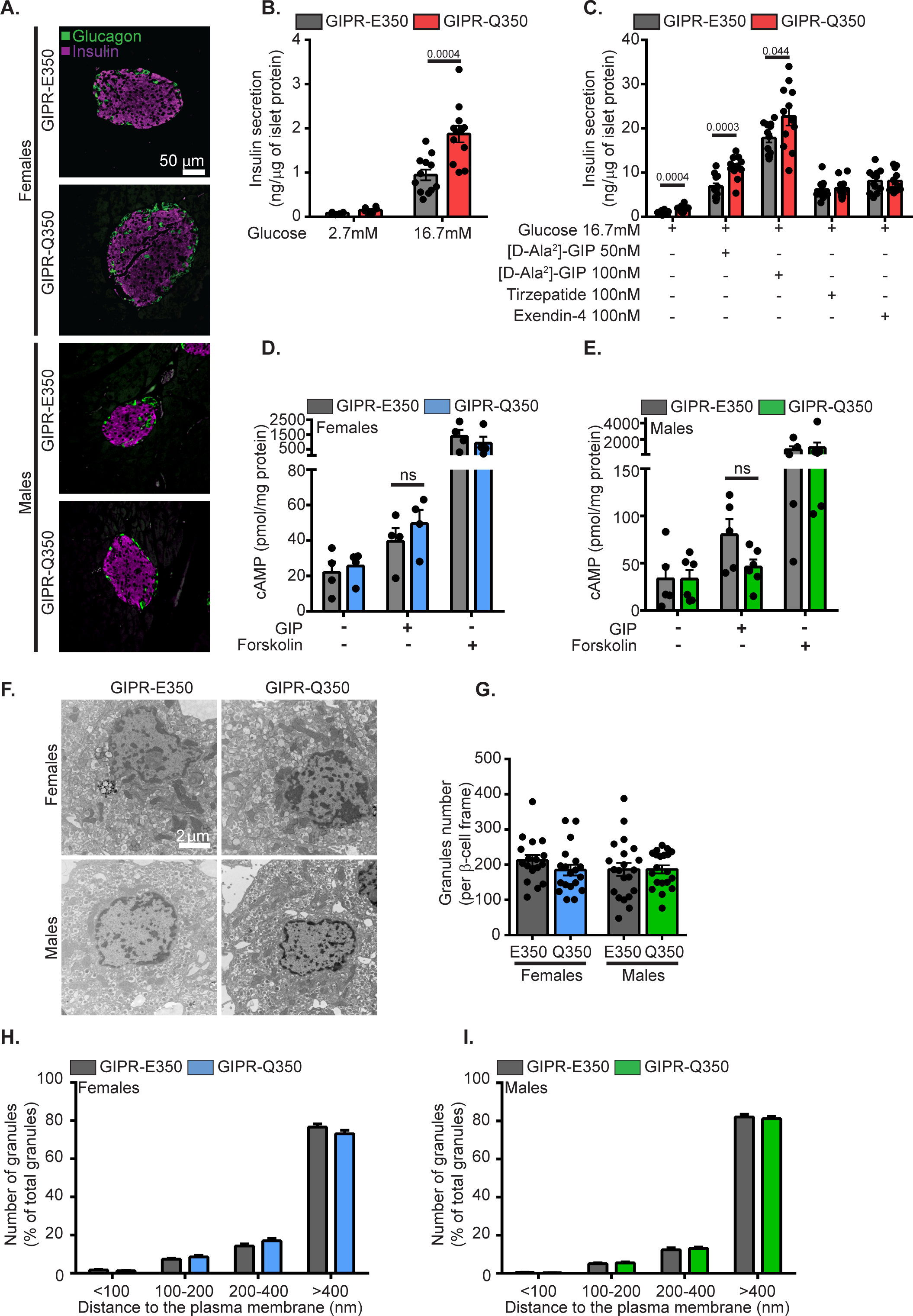
Insulin secretion is increased in GIPR-Q350 islets. **A.** Immunofluorescence of pancreatic islets. Staining shows Glucagon (green) and Insulin (magenta). Scale bar, 50 μm. **B&C**. GSIS performed on 8 to 10 technical replicates per condition of pooled islets of similar sizes isolated from GIPR-E350 and GIPR-Q350 mice (n = 3 mice/genotype). Islets were incubated for 45 min in 2.7 mM or 16.7 mM Glucose **(B)**, or in 16.7 mM Glucose with or without [D-Ala^2^]-GIP (50 nM or 100 nM), Tirzepatide (100 nM), Exendin-4 (100 nM) **(C)**. Graphs show Insulin secretion corrected to islets protein content (representative graphs of n = 3 independent experiments). **D&E.** cAMP production was measured on 3 technical replicates per condition of pooled islets of similar sizes isolated from GIPR-E350 (n = 3 – 5) and GIPR-Q350 (n = 3 – 5) female **(D)** and male **(E)** mice. Islets were incubated for 30 min in 16.7 mM Glucose with or without GIP (100 nM) or Forskolin (10 μM). Graph shows cAMP production corrected to total protein content. **F.** Electron microscopy images of pancreatic islets isolated from GIPR-E350 and GIPR-Q350 animals at 22 to 26 weeks of age. Scale bar, 2 μm. **G.** Granule density per β-cell frame in GIPR-E350 and GIPR-Q350 mice (2 to 5 β-cell frame per animal, 5 mice per group). **H&I**. Distance of insulin granule center to the plasma membrane was measured in GIPR-E350 and GIPR-Q350 female **(H)** and male **(I)** mice. Insulin granules were grouped with respect to their distance to the plasma membrane. (2 to 5 β-cell frame per animal, 5 mice per group). Data are mean ±SEM. Two tailed unpaired *t*-tests for B-E.

We then isolated islets from GIPR-E350 and Q350 mice to assess the direct effect of GIP on GSIS. In agreement with the *in vivo* data, GIPR-Q350 islets had an enhanced GSIS compared to GIPR-E350 islets (**Figure 5B**). While [D-Ala^2^]-GIP enhanced GSIS from islets of both genotypes in a dose dependent manner, the incretin effect was significantly more pronounced in GIPR-Q350 islets (**Figure 5C**). This effect was restricted to [D-Ala^2^]-GIP, as Tirzepatide (GIPR and GLP-1R agonist [38; 39]) and Exendin-4 (GLP-1R agonist) treatments did not result in significant changes in insulin secretion between genotypes (**Figure 5C**). There were no significant differences in insulin secretion observed at a 2.7 mM glucose concentration regardless of treatments (**Fig. S5E**)

These *ex vivo* data highlight two functional cell-intrinsic differences between GIPR-E350 and GIPR-Q350 islets. First, the increased GSIS recapitulates the *in vivo* phenotypes and thereby confirms that GIPR-Q350 islets release more insulin when stimulated with glucose than GIPR-E350 islets. Second, increased insulin secretion from GIPR-Q350 islets upon co-stimulation with GIP and glucose establishes an enhanced acute response to GIP.

Next, we sought to elucidate the mechanism by which the GIPR-Q350 variant improves GIP sensitivity in the isolated islets. GIPR signals through Gs leading to cAMP elevation [40; 41]. Consistent with previous studies in cultured cells [25; 34], we found no increase in cAMP production following GIP stimulation (**Figures 5D,E**). Insulin secretion from β-cells is achieved by regulation of granule recruitment to and fusion with the plasma membrane [42; 43], and we did not identify qualitative differences in granule morphology nor differences in the number of insulin granules that would account for the differences in GSIS or GIP incretin effect between the genotypes (**Figures 5F,G**). Furthermore, there were no differences in the distribution of distances of granules to the β-cell plasma membrane between GIPR-E350 and GIPR-Q350 mice (**Figures 5H,I**).

### GIPR-Q354 trafficking is altered in MIN6 β−cells

Previous studies have shown differences in post-activation trafficking between GIPR-Q354 and GIPR-E354 in adipocytes [25; 26]. Post-activation GIPR-Q354 accumulates in the TGN, significantly delaying its return to the plasma membrane as compared to GIPR-E354. We studied the behavior of the two GIPR variants in MIN6 cells, a murine cultured β cell line, using the previously characterized HA-GIPR-GFP constructs [25]. GIP stimulation resulted in a 30% downregulation of plasma membrane GIPR-E354 (**Figure 6A**). In unstimulated cells, GIPR-Q354 expression at the plasma membrane was comparable to that of GIPR-E354; however, GIP-stimulated downregulation of GIPR-Q354 was significantly enhanced, demonstrating differences in post-activation trafficking between GIPR-E354 and GIPR-Q354. As we have previously shown in adipocytes, both GIPR-E354 and GIPR-Q354, following activation, traffic through the TGN (**Figure 6B**).

**Figure 6:**
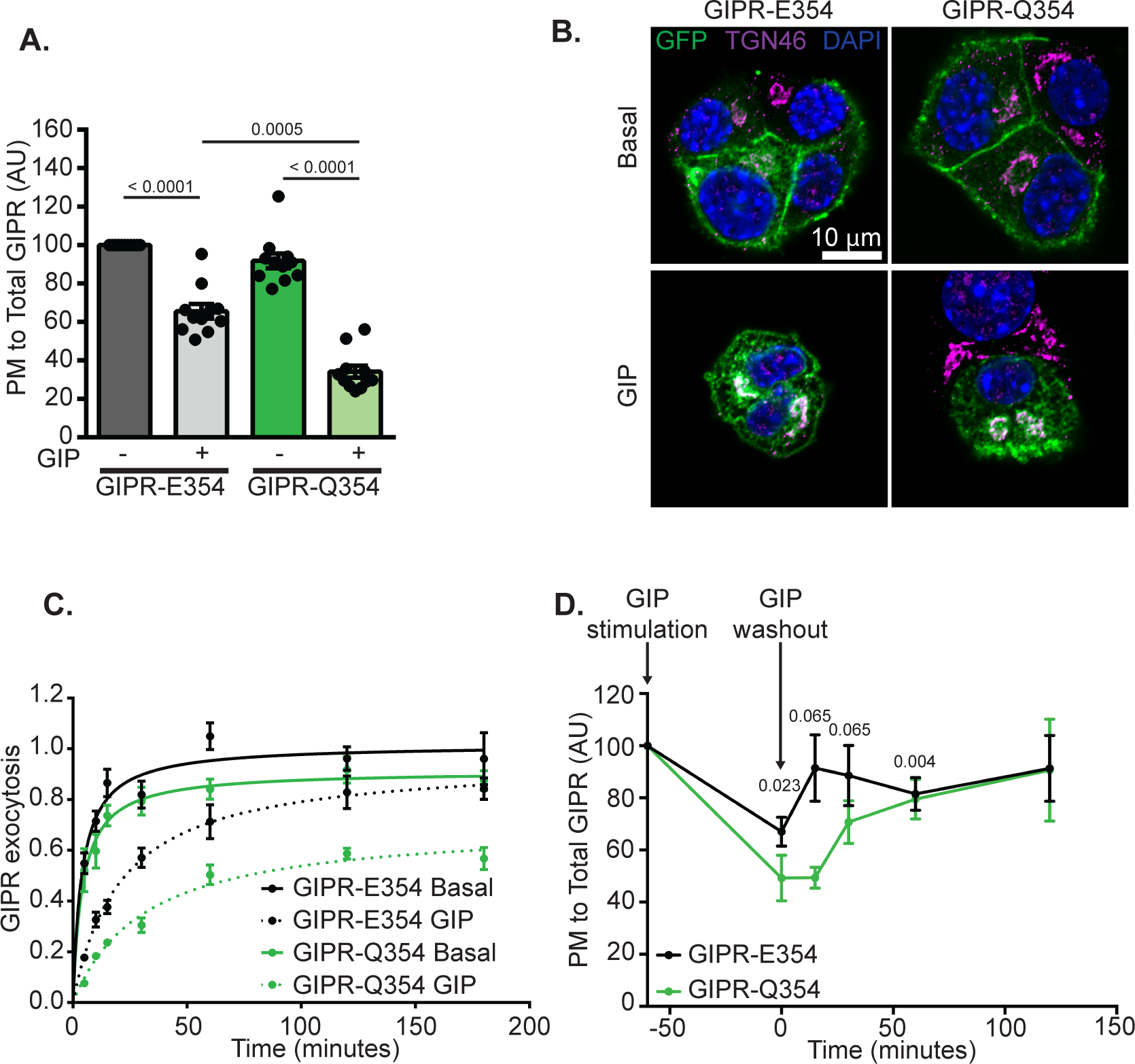
Post activation recycling of GIPR-Q354 is impaired in β−cells. MIN6 cells were electroporated with HA-GIPR-E354-GFP or HA-GIPR-Q354-GFP **A.** Quantification of GIPR plasma membrane (PM) level to total distribution in basal and GIP-stimulated (100nM) cells for 1 h. Data from individual experiments are normalized to the HA-GIPR-E354-GFP-electroporated cells in basal condition (n = 11 independent experiments). **B.** Cells were stimulated or not with GIP (100 nM) for 1 h. Immunofluorescence shows HA-GIPR-E354-GFP or HA-GIPR-Q354-GFP (green), TGN46 (magenta) and nuclei counterstained with DAPI (blue). Scale bar, 10 μm. **C.** Cells were stimulated or not with GIP (100nM) for 1 h prior to incubation with anti-HA antibodies for the indicated times. Graph shows GIPR exocytosis rate in basal or GIP-stimulated cells (n = 4 independent experiments). **D.** Cells were stimulated with GIP (100nM) for 1 h followed by an up to 2 h washout. Graph shows quantification of GIPR plasma membrane (PM) level to total distribution at different time points (n = 5 independent experiments). Data are mean ±SEM. One tailed paired *t*-tests for A, D.

Downregulation of both GIPR-E354 and GIPR-Q354 resulted from a slowing of activated GIPR recycling back to the plasma membrane (**Figure 6C**). Fitting the data to an exponential rise to a plateau revealed that both GIPR-E354 and GIPR-Q354 were constitutively recycled to the plasma membrane in unstimulated cells at similar rates (recycling rate constants of 0.026 and 0.029 min^−1^, respectively). GIP stimulation induced a near twofold slowing of GIPR-E354 recycling (0.015 min^−1^), whereas the recycling of the GIPR-Q354 was reduced fourfold (0.007 min^−1^). In addition, about 30% of the internalized activated GIPR-Q354 did not recycle or did so very slowly, reflected in the reduced plateau level for GIP-stimulated GIPR-Q354 (**Figure 6C**). These data demonstrate that both GIPR-E354 and GIPR-Q354 constitutively recycle in unstimulated MIN6 β−cells, and that GIP stimulation slows the recycling of both GIPR variants, albeit with a significantly more pronounced effect on GIPR-Q354. These findings are in agreement with previous studies of GIPR in cultured adipocytes [25; 26].

An additional consequence of the enhanced downregulation of GIPR-Q354 is that when GIP stimulation was terminated, repopulation of the plasma membrane with GIPR-Q354 takes about four times longer than for GIPR-E354 (**Figure 6D**). However, the plasma membrane levels of GIPR-Q354 are ultimately restored to the pre-stimulus level upon termination of GIPR stimulation, albeit at a slower rate. Thus, the approximate 30% of GIPR-Q354 that does not recycle in stimulated cells (reduced plateau **Figure 6C**) does recycle upon termination of GIPR stimulation (**Figure 6D**).

In sum, these analyses demonstrate that in MIN6 β-cells, as is the case in adipocytes, there are significant differences between the post-activation trafficking of GIPR-E354 and the GIPR-Q354 variant associated with metabolic alterations in humans. Those differences could play a role in the phenotypes observed in our animal models.

## Discussion

GWAS have identified GIPR variants linked to T2D and obesity [27; 44-47]. Here we generated an isogenic mouse model of the naturally occurring human GIPR-Q354 variant to isolate the effect of the variant on whole body metabolic control. In line with the GWAS, the Q350 substitution impacts metabolic tone of mice.

GIPR-Q350 mice of both sexes have an increased sensitivity to GIP to induce glucose clearance when tested by an intraperitoneal GIP + glucose tolerance test. This enhanced glucose clearance in GIPR-Q350 mice was paralleled by increased 15 min circulating insulin, revealing a role for β-cells in the enhanced response. The increased sensitivity is specific to GIP because response to GLP-1 was not altered in GIPR-Q350 variant mice. In addition, when considered across multiple cohorts, GIPR-Q350 mice were more glucose tolerant than GIPR-E350 mice in an intraperitoneal glucose tolerance test. This increased tolerance, although variable, was on the order of a 9 to 12% reduction in the AUC (males and females, respectively), similar in magnitude to the reduction in AUC achieved in GIPR-E350 by GIP (that is, the incretin effect).

We exploited the GIPR-Q350 mice to interrogate alterations in the response of β-cells to glucose and incretin hormones. There were no differences in islet size, morphology, β-cell mass or in the expression of *Gipr*, *Glp1r*, *Glut2* and *Glucokinase* at the mRNA level. Nonetheless, in agreement with the *in vivo* data, GIPR-Q350 islets had enhanced GSIS and increased sensitivity to GIP as compared to GIPR-E350 islets. The increased GIP sensitivity of GIPR-Q350 islets reflects a difference between genotypes in acute response to GIP. As is the case *in vivo*, there is no difference between genotypes in response to GLP-1.

We propose a spatiotemporal difference in signal transduction, specifically the increase dwell time in the TGN of the Q variant, underlies the physiologic differences between GIPR-Q350/4 and GIPR-E350/4 in mice and humans. GPCR signal transduction is not limited to receptor activation at the plasma membrane, but internalized receptors continue to signal, and the post-activation traffic of GPCRs is a critical parameter in sculpting signal transduction [48; 49]. We have previously described differences in post-activation trafficking of GIPR-Q354 compared to GIPR-E354 in studies of cultured adipocytes [25; 26]. A key difference is that the GIPR-Q354 variant is more slowly recycled back to the cell surface following GIP stimulation than is GIPR-E354. This slower recycling results in enhanced accumulation of GIPR-Q354 in the TGN. Here we confirm similar differences in post-activation trafficking of GIPR-Q354 and GIPR-E354 in β-cells, establishing that the differences in trafficking are intrinsic to the GIPR allele and not a result of the cell type used for study. A recent study in cultured β-cells provides additional evidence highlighting the importance of post-activation trafficking of incretin receptors [50]. Following activation by their ligands, the GLP-1R was more efficiently targeted for degradation, whereas GIPR was more efficiently recycled back to the plasma membrane, a difference that might underlie differences in signal transduction between the two incretin receptors.

The specific pathways linking GIP (or GLP-1) to enhanced insulin secretion have not been described in complete molecular detail. GIP enhancement of GSIS is dependent on elevated cAMP produced downstream of GIPR activation [19]. cAMP activation of protein kinase A (PKA) and cAMP-activated guanine nucleotide exchange factor/exchange proteins (EPACs) contribute to GIP augmentation of insulin secretion, although how these signals intersect with insulin granulate release is not understood [51–55]. Despite cAMP being key for GIP signal transduction, we show in primary islets, as has been shown previously in cultured cells [25; 26; 33; 34], that there are no differences in total cAMP production between GIPR-E350 and GIPR-Q350. However, cAMP signaling is spatially restricted [56; 57] and therefore the site of cAMP production might impact signaling downstream of GIPR, differences which might not be captured by whole cell cAMP measurements.

The increased glucose response of GIPR-Q350 islets, independent of GIP stimulation, supports the hypothesis that GIPR-Q350 β-cells, over time, adapt to the GIP hypersensitivity by altering the expression of key element(s) of the glucose-responsive machinery in β-cells. There is evidence that in addition to acute effects on insulin granule release, GIP also effects transcriptional programs, specifically those associated with survival and anti-apoptosis [58; 59]. Transit through the TGN is linked to transcriptional regulation downstream of GPCRs [60–63]; therefore, the increased localization of the Q-variant in the TGN could provide enhanced access to transcriptional signaling responsible for increased GSIS characteristic of GIPR-Q350 β-cells.

When assessed *ex vivo* by GSIS, there is no difference between genotypes in response to Tirzepatide, an artificial hormone comprised of GIP and GLP-1 sequences [17; 38; 39; 64]. Previous reports have identified a biased engagement of Tirzepatide in favor of GIPR trafficking, reflected by Tirzepatide promoting downregulation of the GIPR to the same degree as GIP, whereas it is less effective than GLP-1 at inducing downregulation of GLP-1R [65]. Here we found the GIPR-Q350 islets treated with Tirzepatide secreted insulin to the same amplitude as Exendin-4 but not to that stimulated by [D-Ala^2^]-GIP. This would suggest that the signaling output achieved by GIP:GIPR-Q354 interaction is not mimicked by Tirzepatide effect on GIPR-Q354.

The improved glucose disposal in an oral glucose challenge of GIPR-Q350 mice is not accompanied by elevated levels of insulin, which was unexpected considering the enhanced insulin secretion when exogenous GIP was tested in IP-GTT and *ex vivo* in GSIS. Rather, in the oral glucose tolerance challenge there was a decrease in native GIP, suggesting reduced GIP secretion to match the increased GIP sensitivity of GIPR-Q350 mice. Importantly, our findings parallel the decreased plasma levels of GIP found in human subjects bearing the GIPR-Q354, in fasting and 2-hour post glucose bolus plasma. [32]. GIPR is widely expressed among peripheral tissues as well as in the brain, and in our model the GIPR-Q350 variant is expressed in all tissues that natively express GIPR. Therefore, our data support feedback regulation linking GIP sensitivity to GIP secretion, without informing on the specific tissues and or mediators involved in the feedback regulation. Of note, plasma insulin has previously been reported to provide the feedback regulation in a study of rats [66]. Thus, it is possible that in the GIPR-Q350 mice the decreased GIP levels are matched to the increased insulin secretion of GIP-stimulated GIPR-Q350 islets. We cannot explain why in the oral glucose challenge the increased glucose tolerance of GIPR-Q350 mice is not coupled to a measurable increase in insulin. It is possible a difference in circulating insulin was not captured in the time points we assayed and or that a difference in GIPR signaling in other tissues (e.g., central effects) contribute to the better glucose tolerance of GIPR-Q350 mice.

GIPR-Q350 female mice are leaner on normal chow diet and GIPR-Q350 males are less susceptible to weight gain on a high fat diet, results that agree with GWAS studies identifying GIPR-E354 major allele as a risk factor for obesity [30; 31]. Our results therefore link hypersensitivity to GIP with reduced weight, which contrasts with GIPR whole body knockout mice that are resistant to diet-induced obesity [67; 68]. We do not know how to reconcile the differences between the GIPR-Q/E mice and the GIPR whole body knockouts. In our model the GIPR-Q350 variant is expressed in all tissues that normally express GIPR; therefore, the metabolic phenotype of the GIPR-Q350 variant reflects the sum of the effects on all these tissues, and there is evidence associating GIPR activity in specific tissues to the control of body weight. In peripheral tissues, restoring GIPR expression in white adipose tissue of whole-body GIPR knockout mice is sufficient to restore normal weight gain on a HFD, and adipose-specific knockout of GIPR protects against diet-induced obesity [69; 70]. In contrast, β-cell-specific ablation of GIPR does not prevent against diet-induced obesity [71]. Centrally, GIPR is expressed in the hypothalamic nuclei, a region of the central nervous system (**CNS**) responsible for energy expenditure and food intake [72]. CNS-GIPR KO mice are resistant to diet induced obesity and have an improved metabolism [73]. An additional complication to the interpretation of the knockout phenotype is the more recent appreciation of GIPR antagonism by cleaved forms of GIP, in addition to GIPR agonism, and how either contribute to metabolism modulation [74].

Pharmacological agonism of incretin hormones has been proposed as a treatment of T2D. GLP-1 receptor agonists were shown to impact satiety, induce weight loss, enhance GSIS and regulate glucose homeostasis [75; 76]. In GIPR based therapy, there is a dichotomy between agonism and antagonism of the receptor [77]. Studies have targeted either GIP [68; 78] or GIPR [79] with antagonist antibodies and have revealed a protective effect against diet-induced body weight gain. Others have used GIPR peptide agonists to show that by chronically infusing either GIP analog [80] or by a dual treatment with a GLP-1 analog [81], a decreased weight and metabolic benefits can be achieved. These discrepancies argue that a modulation rather than a complete inhibition or chronic activation of GIPR activity is key to a better metabolism. Here we link spatiotemporal differences in intracellular trafficking of GIPR with metabolic alterations, opening a potential new direction for targeting the trafficking of GIP receptors to modulate receptor signaling controlled by native GIP.

### Limitations of the study

The model presented describes the phenotypes and provides some mechanistical understanding of how GIPR-Q354 functions. This study demonstrates the importance of intracellular trafficking of GPCRs and its impact on whole body metabolism. While exact mechanisms need further investigation, this work sets the basis for generation of new hypothesis on the mechanisms underlying GIPR role on glucose homeostasis and T2D. However, because inter-organ communication is critical in the homeostatic regulation of metabolism, our studies focusing on differences in β-cells may only account for some of the differences between the genotypes.

In PheWAS analyses, human subjects bearing the GIPR-Q354 SNP have a lower BMI and paradoxically an increased risk of T2D as compared to GIPR-E354 subjects. In our mouse model, we recapitulate the anthropometric traits observed in humans as well as identify metabolic differences between the GIPR genotypes. However, the GIPR-Q350 variant in our mouse model is not associated with insulin-resistance, rather the GIPR-Q350 mice are more glucose tolerant and sensitive to GIP. Thus, our work provides the basis for a new direction in incretin biology research focused on understanding how increased sensitivity to GIP might lead to an increased risk of T2D in humans harboring the GIPR-Q354.

## Acknowledgments

We thank Marissa Cortopassi, Dr. Hayley Nicholls and Dr. David Cohen for their assistance with Promethion metabolic cages experiments. We thank the Weill Cornell Medicine Metabolic Phenotyping Center, the Weill Cornell Medicine Biochemistry Microscopy & Image Analysis Core and the Human Islet and Adenovirus Core of the Einstein-Sinai Diabetes Research Center (NIH P30 DK020541). We thank David Soares for technical assistance and other members of the McGraw lab.

This work was supported by NIH grants R01 DK096925 (T.E.M.), R01 DK125699 (T.E.M.), and R01 DK121140 (J.C.L), and by American Diabetes Association Postdoctoral Fellowship grant 1-19-PMF-026 (B.P.)

## Author Contributions

L.Y. designed and conducted experiments, analyzed the data, prepared figures, and wrote the manuscript. B.P. conducted experiments, analyzed the data, and edited the manuscript. N.A. designed and conducted experiments and analyzed the data. R.A.L. and E.F.J. conducted experiments and analyzed the data. N.G-B. and C.R. performed some GSIS experiments. J.W. performed *in vitro* experiments. T.H assisted with animal studies. M.D.G analyzed the indirect calorimetry data. J.C.L. and A.G-O. supervised some experiments. T.E.M. conceived and supervised the project, designed experiments, analyzed the data, and wrote the manuscript.

## Methods

### Experimental models

#### Animals

C57BL/6J mice were purchased from Jackson Laboratory. The mice were generated in the WCM MSKCC animal facility. We generated C57BL/6J GIPR-Q350 variant mice using CRISPR-CAS9 editing of ES cells, the equivalent of the GIPR-Q354 variant in human. We designed the sgRNA 5’- AATCAGGGGCGAGGACATGCGGGCCACTGACATCCGTCTCTTCCCAGGCTGGCTCGCTC CACGCTGACACTGGTGCCCCTGCTGGGTGT**A**CACGT**C**GTGTTTGCGCCTGTGACGGAGG AACAGGTTGAAGGCTCCCTGCGCTTCGCCAAACTGGCCTTTGAAATCTTCCTAAGTTCCTTCCAGGTGC-3’ targeting the exon 11 on the GIPR gene, to introduce a point mutation in the GIPR sequence, as well as restriction sites to accurately genotype the founders. Genotyping was subsequently confirmed by PCR and by sequencing of 1000 bp surrounding the mutation using the following primers: forward 5’- TAAGGTGAGGGCAGGCTCAGGAC-3’ and reverse 5’- CTCCAGTCAATAGTGGCTACTCTG-3’ (**Appendix 1**).

**Appendix 1:**
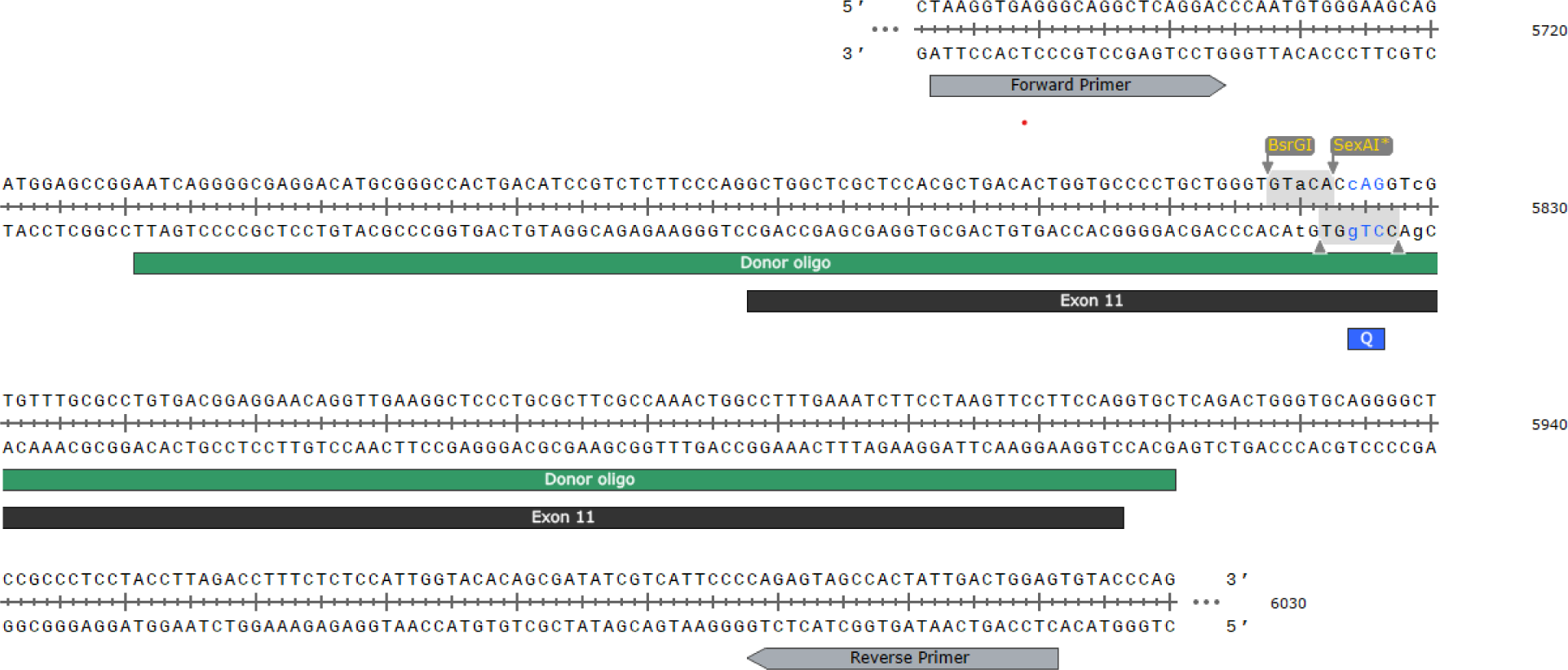
Description of the targeting strategy for GIPR-Q350 mice generation. Created with SnapGene®

Once founders were identified, mice were further bred back to C57BL/6J mice for 10 generations after which male and female mice homozygous for Q350 allele were used for studies. Over the course of this study 13 cohorts of mice have been analyzed. All experiments were conducted on mice between 8 to 25 weeks of age, except for the weights and glucose over time measurements experiments in Figures 2A-D.

Mice were maintained on a 12 h/12 h light/dark cycle at room temperature and had ad libitum access to food and water. Mice were fed a chow diet (5053, PicoLab® Rodent Diet), except for the male mice study on high fat diet where mice were fed a 60% high fat diet (D12492i, Research Diets).

The Institutional Animal Care and Use Committee and the Research Animal Resource Center at Weill Cornell Medical College approved all animal procedures.

#### Cell lines

MIN6 cells were grown in DMEM (12100046, ThermoFisher Scientific) supplemented with 15% Fetal bovine serum (FBS, 26140-095 ThermoFisher Scientific), 1 % Penicillin-Streptomycin (15070063, ThermoFisher Scientific), 2 mM Glutamine (25030-081, ThermoFisher Scientific), 50 μM β-mercaptoethanol (63689, Sigma-Aldrich) and kept at 37°C and 5% CO_2_.

### Method details

#### cDNA constructs and electroporation

MIN6 cells were electroporated with 45 μg of either HA-GIPR-E354-GFP or HA-GIPR-Q354-GFP plasmids and cultured on coverslips. Experiments were conducted 24 h after electroporation. Cells were serum starved prior to incubation with GIP (100 nM, H-3824.0500, Bachem) for the indicated times.

#### Quantification of Surface to total GIPR

Electroporated MIN6 cells cultured on coverslips were serum starved and treated with or without GIP (100 nM). Cells were fixed and stained with anti-HA antibodies (901503, BioLegend) without permeabilization. After PBS washes, cells were incubated with anti-mouse Cy3-conjugated antibodies. Cells were imaged by epifluorescence using a 20x objective (Leica Biosystems) and the intensity ratio of Cy3/GFP was used as an indicator of surface GIPR/total GIPR. Intensity for each cell was quantified using MetaMorph software.

#### Quantification of GIPR exocytosis

Electroporated MIN6 cells cultured on coverslips were serum starved and treated with or without GIP (100 nM) for 1 h. Live cells were then incubated with anti-HA antibodies for indicated times. Cells were immediately fixed, permeabilized and incubated with anti-mouse Cy3-conjugated antibodies.

#### Metabolic cages

Male mice on a HFD were single-housed and acclimated to the Promethion High-Definition Multiplexed Respirometry Systems (Sable Systems International) at ambient temperature and light controlled environment. Oxygen consumption, carbon dioxide production, food intake, body mass, movement and activity were measured continuously over a period of 24 h. This experiment was conducted through the Weill Cornell Medicine Metabolic Phenotyping Center.

#### Glucose tolerance test

Oral glucose tolerance tests (O-GTT) were performed in 6 h fasted mice. The glucose dose used was 2 g/kg.

Intraperitoneal glucose tolerance tests (IP-GTT) were performed in 16 h fasted mice. Mice were either ip injected with glucose at a dose of 2 g/kg or with glucose 2 g/kg and GIP 20 μmol/kg. For [D-Ala^2^]-GIP (6699, Tocris), mice were injected with a single dose of [D-Ala^2^]-GIP (0.1 μg, 0.5 μg, 1 μg) 30 min prior to glucose 2 g/kg injections. For Exendin-4 (E7144, Sigma Aldrich), mice were injected with a single dose of Exendin-4 (1 μg) 30 min prior to glucose 2 g/kg injections.

For both O-GTT and IP-GTT, blood glucose level was measured at indicated time points and blood was collected at 0, 7 and 15 min in capillary microvettes coated with K3 EDTA for O-GTT and 0 and 15 min for IP-GTT and centrifuged for 15 min at 4°C to collect plasma.

#### Insulin tolerance test

Insulin tolerance tests (ITT) were performed in 6 h fasted mice. Mice were intraperitoneally injected with 0.75 U/kg Insulin. When co-injection with GIP 20 μmol/kg was performed during the ITT, male mice were injected with 0.75 U/kg Insulin and females 0.5 U/kg Insulin.

#### Islets isolation

Pancreata were perfused with Collagenase P (1.7 mg/ml, 11249002001, Sigma-Aldrich) via the pancreatic duct. Pancreata were collected, digested at 37°C for 15 min and washed in HBSS supplemented with 10% FBS. Islets were isolated using Histopaque® (10771, Sigma-Aldrich) gradient. Islets were handpicked and allowed to recover overnight in RPMI (11879-020, ThermoFisher Scientific) media supplemented with 10% FBS, 1 % Penicillin-Streptomycin and 5.5 mM Glucose.

#### Glucose stimulated insulin secretion

For each experiment, islets from 3 mice per genotype were pooled and incubated in basal KREBS media (119 mM NaCl, 100 mM HEPES, 23 mM KCl, 5 mM MgSO_4_, 0.75 mM Na_2_HPO_4_, 2 mM KH_2_PO_4_, 25 mM NaHCO_3_, 2mM CaCl_2_, 0.05% BSA) supplemented with 2.7 mM Glucose for 2 h at 37°C and 5% CO_2_. 5-10 replicates of 5 islets per genotype were incubated in 2.7 mM Glucose or 16.7 mM Glucose with or without [D-Ala^2^]-GIP (50 nM and 100 nM), Tirzepatide (100 nM) (Peptide Sciences), or Exendin-4 (100 nM), and in the presence of 200 μM IBMX for 45 min at 37°C and 5% CO_2_. Supernatants were collected and immediately put on ice. Islets were lysed in RIPA buffer with anti-phosphatase and anti-protease inhibitors and were used to measure total protein content. Supernatants and protein lysates were stored at −20°C until Insulin ELISA was performed.

#### cAMP production

Islets from 3-5 mice per genotype were pooled and incubated in basal KREBS media supplemented with 2.7 mM Glucose for 2 h at 37°C and 5% CO_2_. 3 replicates of 25 islets per genotype were incubated with 16.7 mM Glucose with or without GIP (100 nM) or Forskolin (10 μM) (F3917, Sigma Aldrich) in the presence of 200 μM IBMX for 30 min at 37°C and 5% CO_2_. Islets were lysed in 1N HCl, sonicated at 37 kHz for 45 sec. cAMP ELISA was performed on lysates.

#### Quantitative RT-PCR

24 h after islets isolation, islets were lysed and RNA was extracted using the RNeasy kit (74106, Qiagen), according to the manufacturer’s protocol. cDNA was obtained using the RNA to cDNA EcoDry Premix (639545, Takara). Quantitative PCR was performed with iQ SYBR Green Supermix (170-8884, Bio-Rad). Oligonucleotides used in this study: Hprt 5’- TGCTCGAGATGTCATGAAGG-3’ (Forward) and 5’- TATGTCCCCCGTTGACTGAT-3’ (Reverse); Gipr 5’- AAAGATGTTGGAGACCACAGAAC-3’ (Forward) and 5’- GCAGACACCTGACGGAACC-3’ (Reverse); Glut2 5’- GGGACAAACTTGGAAGGATCAA-3’ (Forward) and 5’-AAATTTGGAACATCCCATCAAGAG-3’ (Reverse); Glucokinase 5’- TGAGCCGGATGCAGAAGGA-3’ (Forward) and 5’- GCAACATCTTTACACTGGCCT-3’ (Reverse); Glp1r 5’- ACGGTGTCCCTCTCAGAGAC-3’ (Forward) and 5’- ATCAAAGGTCCGGTTGCAGAA-3’ (Reverse).

#### Immunofluorescence

MIN6 cells were cultured on coverslips and fixed with 3.7% formaldehyde (F1635, Sigma-Aldrich). Cells were treated with 0.1% Triton X-100 for 10 min. Nonspecific binding was blocked by incubating cells in blocking buffer (10% calf serum in PBS) for 30 min. Incubation with primary antibodies was performed at 37°C for 1 h and incubation with secondary antibodies was performed at 37°C for 30 min.

7 μm paraffin embedded mice pancreas sections were antigen retrieved in Citrate pH 6.0 and blocked with 3% BSA, 10% FBS in PBS buffer for 30 min. Sections were incubated with primary antibodies in blocking buffer overnight at 4°C prior to incubation with Alexa 488 and Cy3-conjugated secondary antibodies for 30 min at room temperature.

Antibodies used for immunofluorescence: Mouse monoclonal anti-Insulin (I2018, Sigma-Aldrich), Rabbit monoclonal anti-Glucagon (8233, Cell Signaling Technology), Rabbit polyclonal anti-TGN46 (ab16059, Abcam), Mouse monoclonal anti-HA (901503, BioLegend), Goat polyclonal anti-mouse Cy3 (115-165-062, Jackson Immunoresearch), Donkey polyclonal anti-rabbit Alexa Fluor™ 488 (A-21206, Thermofisher Scientific), Horse polyclonal anti-mouse IgG (H+L), biotinylated (BA-2000, Vector Laboratories)

Images were acquired using a Zeiss LSM 880 with Airyscan confocal microscope and prepared using ImageJ software.

#### Immunohistochemistry for beta cell mass analysis

8 serial 5 μm paraffin embedded mice pancreas sections separated by 100 μm per animal were antigen retrieved in Citrate pH 6.0 and incubated in 0.3% H_2_O_2_ to quench endogenous peroxidases. After Avidin and Biotin blocking, sections were incubated with anti-Insulin antibody diluted in 10% FBS and PBS. The following day, Biotinylated Horse Anti-Mouse IgG Antibody was applied to the sections prior to incubation with VECTASTAIN® Elite® ABC-HRP Reagent. Signal development was obtained using DAB and a hematoxylin counterstaining. Slides were scanned in entirety (Nikon CoolScan) and Pancreatic and islets areas were quantified using ImageJ.

#### Electron Microscopy

Islets from 5 mice per genotype were isolated. Islets were processed at the Weill Cornell Medicine Microscopy and Image Analysis Core Facility and imaged at random using JEOL JEM 1400 Transmission Electron Microscope. The distance from granule center to the plasma membrane was performed using ImageJ and a macro kindly provided by the Office of Collaborative Science (OCS) Microscopy Core at NYU Langone Medical Center.

#### ELISA

Insulin (81527, Crystal Chem), GIP (EZRMGIP-55K, Sigma-Aldrich), and GLP-1 (80-GLP1A-CH0, Alpco) ELISA kits were used to measure their level in the plasma. cAMP ELISA kit (ADI-900-066, Enzo Life Sciences) was used to measure cAMP production in islets. Protocols as provided by manufacturers were followed.

#### Quantification and statistical analysis

All experiments were repeated at least three times. At least 5 mice per condition and per genotype were used as biological replicates for *in vivo* experiments. Results are expressed as means ± SEM. Data were analyzed using Prism software (GraphPad). Groups were compared with an analysis of variance (ANOVA) or a Student’s t test, and a P value < 0.05 was considered as significantly relevant.

**Supplementary Figure 1:**
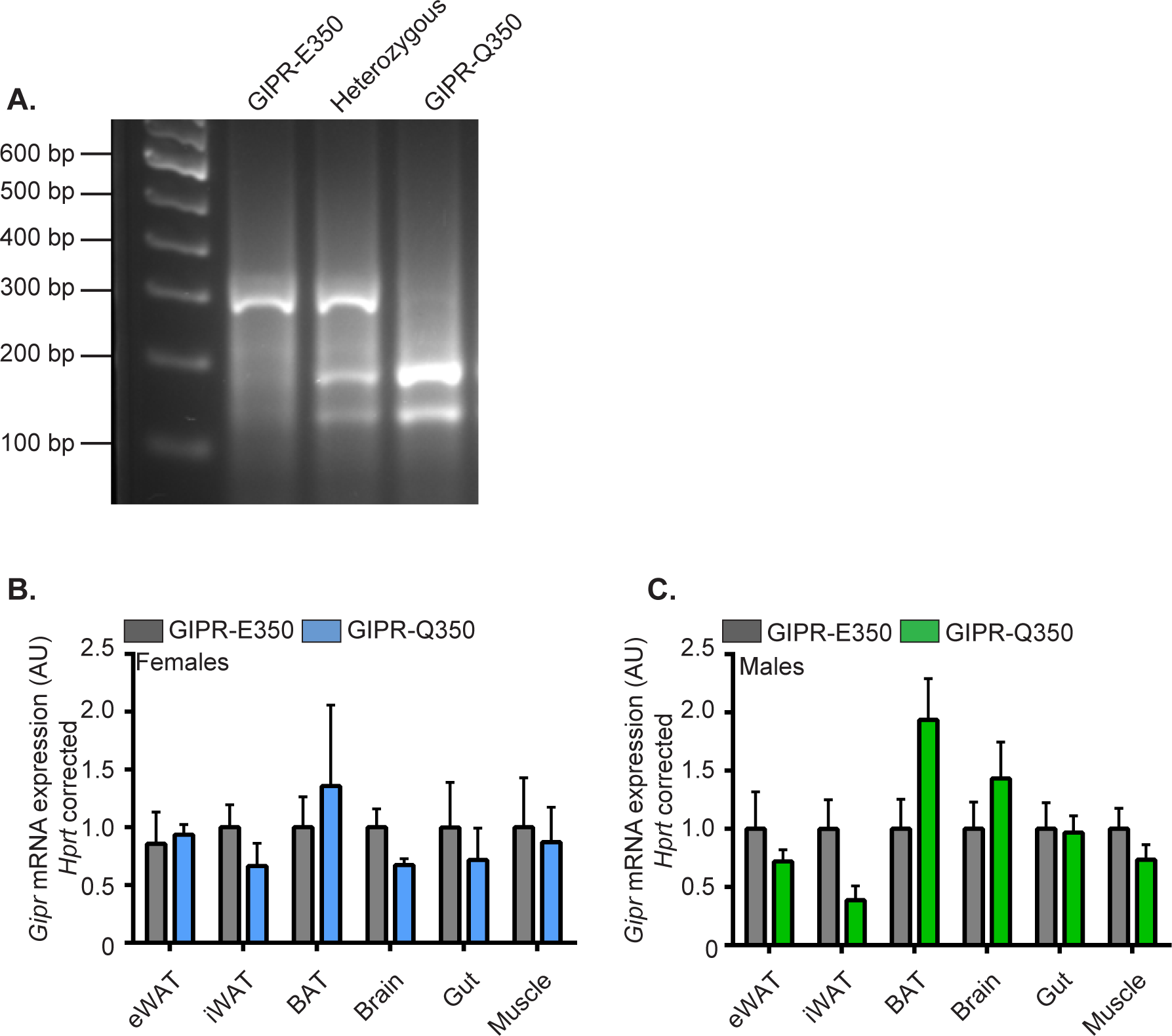
GIPR-Q350 generation and GIPR expression in mice. **A.** Representative PCR gel of GIPR-E350, heterozygous and GIPR-Q350 mice genotyping. **B&C**. Relative mRNA expression of *Gipr* in the epidydimal white adipose tissue (eWAT), inguinal white adipose tissue (iWAT), brown adipose tissue (BAT), brain, gut, and muscle from GIPR-E350 and GIPR-Q350 females **(B)** and males **(C)** (n = 5 – 6 mice/genotype). Data are mean ±SEM.

**Supplementary Figure 2:**
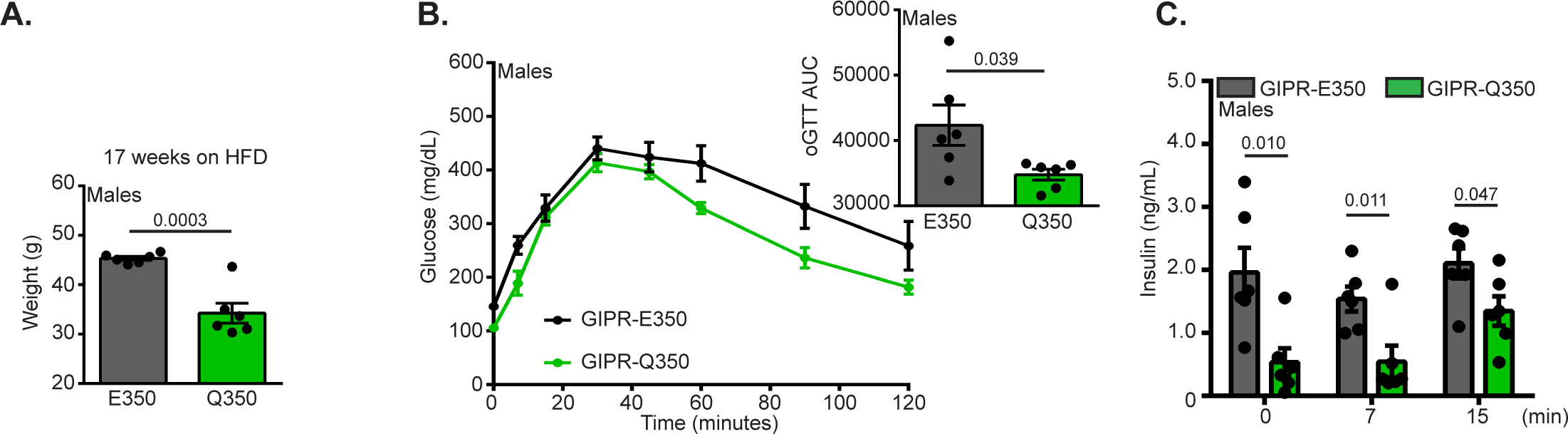
HFD GIPR-Q350 male mice are glucose tolerant. **A.** Weights after 17 weeks of HFD feeding. **B.** Blood glucose excursion and area under the curve (AUC, insert) over time after an oral glucose tolerance test (O-GTT, 2 g/kg) in 16 h fasted mice. **C.** Plasma levels of Insulin before and at the indicated times after glucose administration (2 g/kg). (n = 6 mice/genotype). Data are mean ± SEM. Two tailed unpaired *t*-tests for A-C.

**Supplementary Figure 3:**
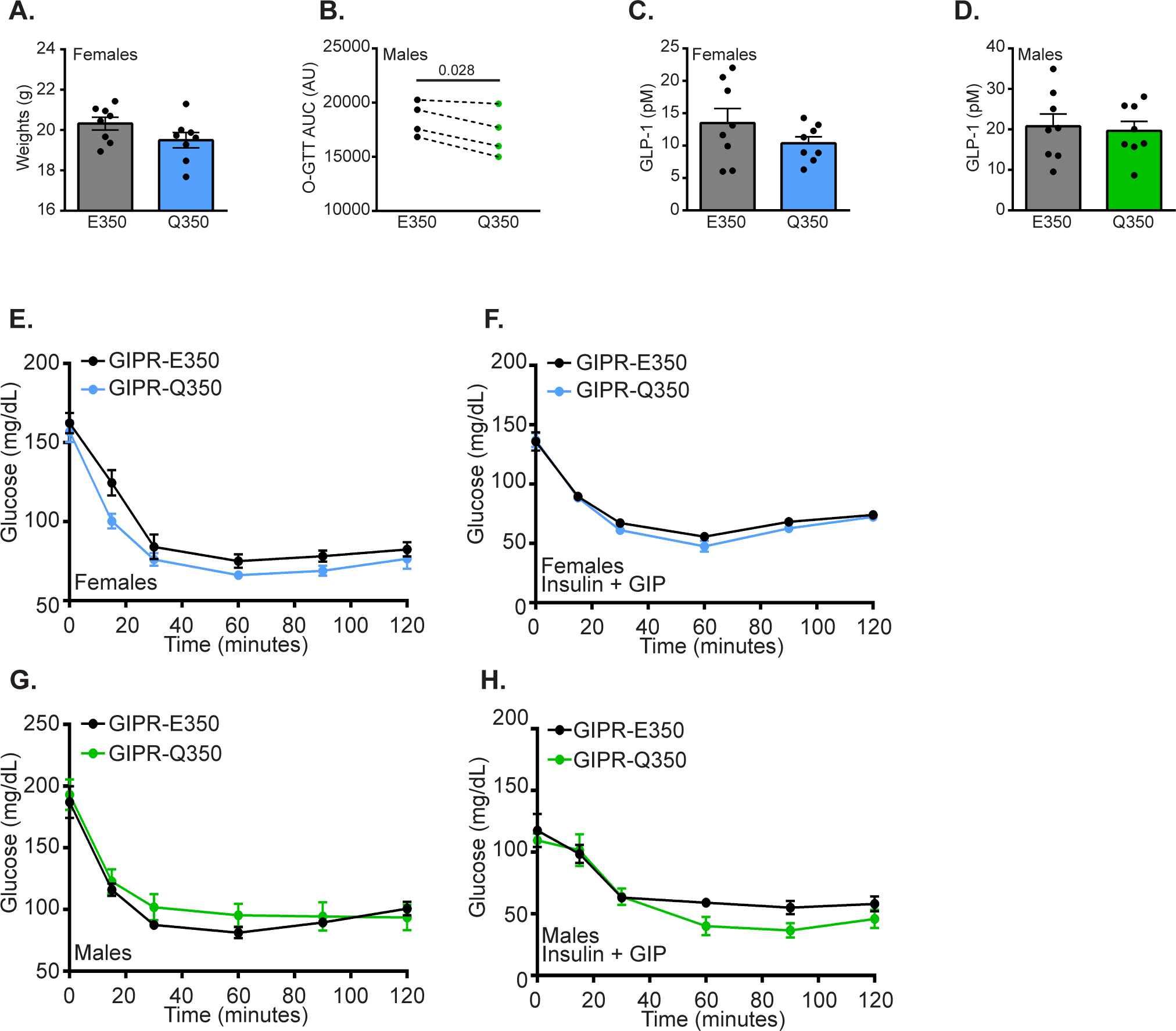
GIPR-Q350 does not impact the insulin sensitivity of mice. **A.** Weights of GIPR-E350 and GIPR-Q350 females prior to an oral glucose tolerance test (O-GTT). (n = 8 mice/genotype). **B.** Mean area under the curve (AUC) after an oral glucose tolerance test (O-GTT, 2 g/kg) in male mice from 4 independent experiments. **C&D.** Plasma levels of GLP-1 15 min after an oral glucose tolerance test (O-GTT, 2 g/kg) in GIPR-E350 and GIPR-Q350 females **(C)** and males **(D)**. (n = 8 mice/genotype). **E.** GIPR-E350 and GIPR-Q350 female mice were fasted for 6 h prior to injection with Insulin (0.75 U/kg) and their blood glucose levels were monitored for 1 h. (n = 18 – 20 mice/genotype). **F.** GIPR-E350 and GIPR-Q350 females were fasted for 6 h prior to co-injection with Insulin (0.5 U/kg) and GIP (20 μmol/kg) and their blood glucose levels were monitored for 2 h. (n = 8 mice/genotype). **H.** GIPR-E350 and GIPR-Q350 male mice were fasted for 6 h prior to injection with Insulin (0.75 U/kg) and their blood glucose levels were monitored for 1 h. (n = 18 – 20 mice/genotype). **H.** GIPR-E350 and GIPR-Q350 males were fasted for 6 h prior to co-injection with Insulin (0.75 U/kg) and GIP (20 μmol/kg) and their blood glucose levels were monitored for 2 h. (n = 8 mice/genotype). Two tailed paired *t*-tests for B.

**Supplementary Figure 4:**
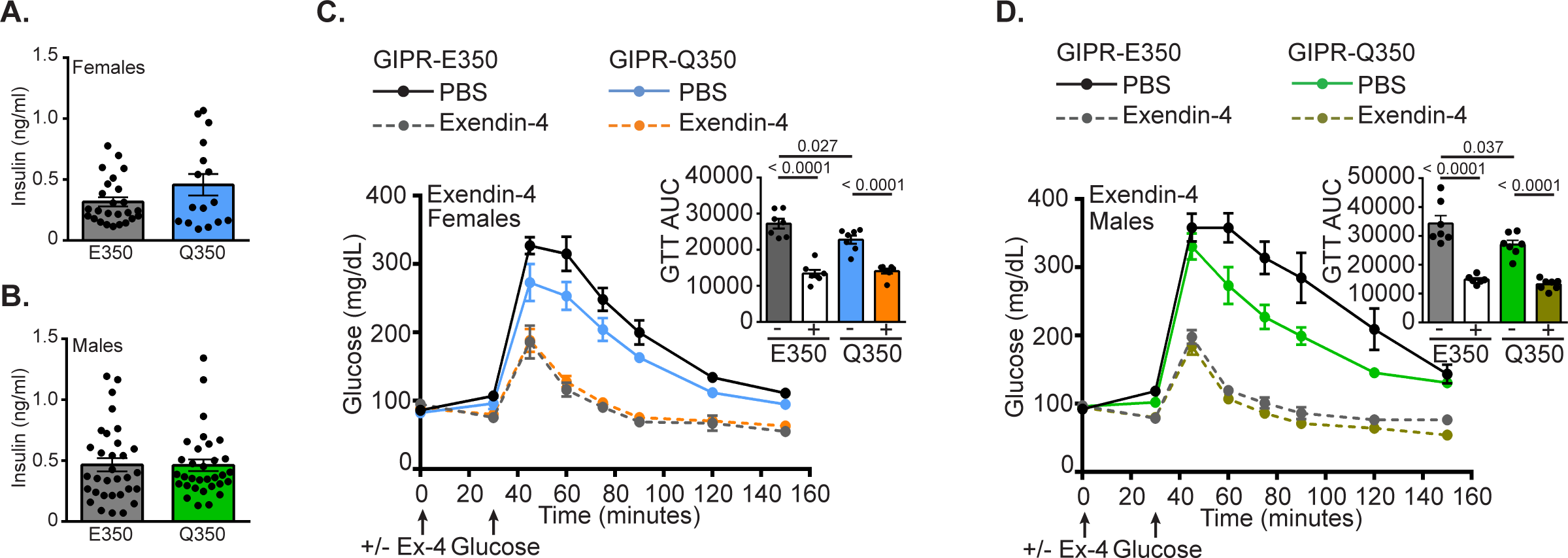
GIPR-E350 and GIPR-Q350 mice have a similar sensitivity to Exendin-4. **A&B**. Plasma levels of Insulin in 16 h fasted GIPR-E350 and GIPR-Q350 females **(A)** and males **(B)**. (n = 16 – 34 mice/genotype). **C&D**. Blood glucose excursion and area under the curve (AUC, inset) after an intraperitoneal glucose challenge (2 g/kg of body weight) assessed 30 min after an i.p. injection of either vehicle (PBS) or Exendin-4 (1 μg) in 16 h fasted GIPR-E350 and GIPR-Q350 female **(C)** and males **(D)** mice. (n = 7 mice/genotype). Two tailed unpaired *t*-tests for C, D.

**Supplementary Figure 5:**
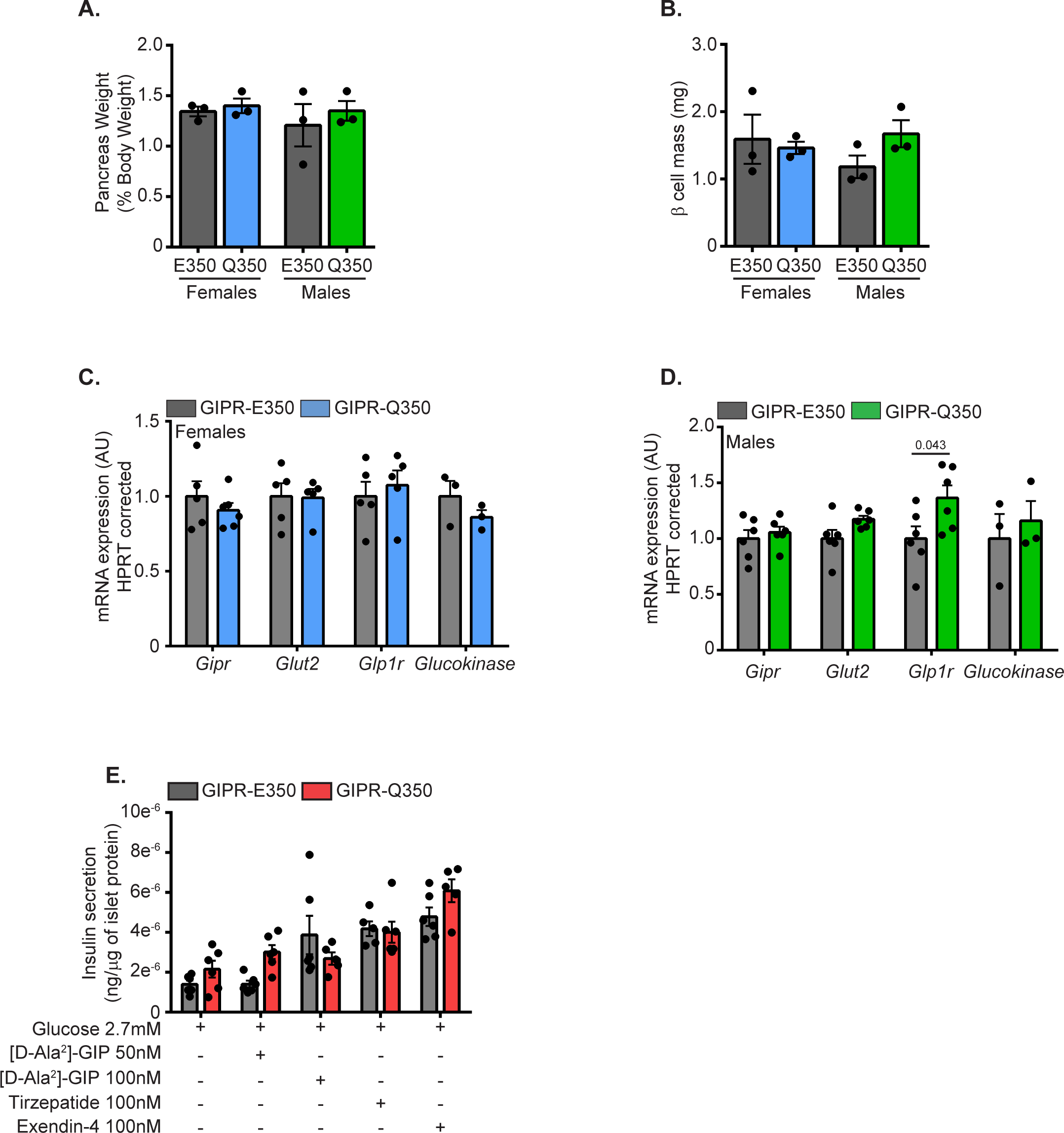
GIPR-Q350 does not impact expression of key metabolic genes in islets. **A.** Pancreas weights corrected to total body weights of GIPR-E350 and GIPR-Q350 mice. **B.** β-cell mass relative to pancreas weights of GIPR-E350 and GIPR-Q350 mice (8 pancreas sections/mouse, n = 3 mice/genotype). **C&D.** Relative mRNA expression of *Gipr*, *Glut2*, *Glp1r* and *Glucokinase* in isolated islets from GIPR-E350 and GIPR-Q350 females **(C)** and males **(D)** (n = 3 – 6 mice/genotype). **E.** GSIS performed on 5 to 6 technical replicates per condition of pooled islets of similar sizes isolated from GIPR-E350 and GIPR-Q350 mice (n = 3 mice/genotype). Islets were incubated for 45 min in 2.7 mM Glucose with or without [D-Ala^2^]-GIP (50 nM or 100 nM), Tirzepatide (100 nM), Exendin-4 (100 nM). Data are mean ± SEM. Two tailed unpaired *t*-tests for D.

